# *Ras^G12V^* Oncogene-Induced Epithelial Senescence and Its Relay Promotes Host Metabolic Syndrome in *Drosophila*

**DOI:** 10.64898/2026.03.13.711573

**Authors:** Saurabh Singh Parihar, Jyoti Tripathi, Sushmita Kundu, Shweta Banerjee, Isha M. Anerao, Pradip Sinha

## Abstract

Precancerous oncogenic activation in a target organ often induces senescence, a tumor-suppressive response known as oncogene-induced senescence (OIS). Clinical observations indicate a strong association of metabolic syndrome (MetS) with the precancerous and early-stage cancers. Notably, cells displaying OIS are characterized by a senescence-associated secretory phenotype (SASP), in which they secrete factors, including inflammatory cytokines. Thus, SASP from cells displaying OIS may trigger host MetS, which likely underpins its association with cancers, such as colorectal cancer (CRC). Here, we tested this hypothesis and show that, in *Drosophila*, the activated *Ras*^*G12V*^ oncogene, which is frequently implicated in human CRC, induces OIS in imaginal disc epithelium and systemically triggers host larval MetS via the conserved cytokine Upd1/IL6. Thus, the larval host with *Ras*^*G12V*^-induced epithelial OIS displays MetS, characterized by obesity, increased lipid and glycogen accumulation in the fat body, and altered insulin signaling, marked by transition from hyperinsulinemia to insulin resistance—all at a precancerous stage. Further, we also noted hyperphagia and increased expression of insulin-like peptides (dILP2/3/5) in the brain of larvae displaying *Ras*^*G12V*^-induced OIS. Notably, *Ras*^*G12V*^-induced OIS is systemically relayed, leading to activation of a senescence-like program in the distant fat body. Genetic suppression of *upd1* or pharmacological intervention with the senomorphic agent, Metformin, attenuated fat body senescence and mitigated MetS-associated phenotypes. Our findings thus identify a causal relationship between OIS and host MetS, suggesting its utility as an early biomarker for detecting cancers such as CRC and its potential as a prophylactic target.

## INTRODUCTION

Multiple clinical and epidemiological studies indicate that metabolic syndrome (MetS)— characterized by obesity, high blood pressure, hyperglycemia, and high cholesterol— markedly increase the risk of developing clinically recognized premalignant precursors such as colorectal adenomas (Kant and Hull, 2011; Zhang et al., 2023b), Barrett’s esophagus (He et al., 2016), and oral premalignancies (Yen et al., 2011). However, MetS and related systemic perturbations—such as the profound fat and muscle wasting seen in cancer-associated cachexia—are classically viewed as secondary outcomes of established cancers, which serve as reservoirs of pro□inflammatory cytokines (Porporato, 2016; Wyart et al., 2025). Yet, this correlation does not establish causality. Whether MetS predisposes to tumorigenesis or instead arises as a metabolic consequence of cancer progression remains unresolved.

Oncogenic lesions add another layer of complexity to this association by inducing cellular senescence, namely, oncogene□induced senescence (OIS) (Hoi et al., 2026; Stangis et al., 2024). OIS acts as an intrinsic, early defense mechanism that delays tumor onset by arresting the proliferation of oncogenically transformed cells (Bartkova et al., 2006; Braig and Schmitt, 2006; Bringold and Serrano, 2000). However, senescent cells secrete pro□inflammatory cytokines, chemokines, and tissue□remodeling factors collectively referred to as the senescence□associated secretory phenotype (SASP) (Coppé et al., 2010; Kumari and Jat, 2021). These SASP factors, mainly pro-inflammatory cytokines, can trigger localized and systemic perturbations, for instance, activation of the paracrine (Acosta et al., 2013; Kuilman et al., 2008; Nelson et al., 2012) and, more recently identified, systemic senescence program (Kiourtis et al., 2024). Thus, these systemic consequences of local senescence may, in turn, promote MetS (Zhang et al., 2023a) and create a physiological milieu that favors cancer progression.

Despite these insights, the role of OIS in shaping the MetS–cancer relationship remains poorly understood. In particular, it is unclear whether preneoplastic senescence can act as a primary driver of systemic MetS, even before overt cancer develops. Here, using a conserved *Drosophila* model of epithelial OIS, we show that discrete preneoplastic lesions can induce local senescence that communicates with distant organs to provoke systemic metabolic dysfunction. Specifically, these precancerous lesions secrete a SASP cocktail rich in the IL□6 homolog Upd1, which signals the brain to trigger hyperphagia and increased insulin production. It also signals the fat body to induce secondary senescence and lipid accumulation—features of MetS. This extended MetS further contributes to adult cachexia.

Our findings establish a causal connection between epithelial OIS and host MetS. We demonstrate that early oncogenic signals can reprogram host metabolism and propagate senescence to distant tissues well before tumor formation. Finally, we show that interrupting this senescence–metabolic axis—either by neutralizing the senescence□relaying cytokine or by treating with the senomorphic and antidiabetic drug Metformin—can suppress the MetS□to□cachexia transition and restore organismal homeostasis.

## RESULTS

### Oncogenic *Ras*^*G12V*^ induces a hyperplastic precancerous lesion with local metabolic reprogramming and OIS-SASP program in the wing imaginal disc

OIS is often observed in precancerous/early-stage cancers (Collado et al., 2005; Hoi et al., 2026). We used a well-characterized mutated version of the *Drosophila* gene *Ras85D (Ras1)*, namely *Ras*^*G12V*^ (Karim and Rubin, 1998), in which valine substitutes glycine at position 12, causing constitutive activation of the RAS GTPase, mimicking its human counterpart responsible for multiple cancers (Bos, 1989; Cox and Der, 2010; Neuman-Silberberg et al., 1984). We used the *vg-Gal4* (Halder et al., 1998; Williams et al., 1994) (vestigial boundary enhancer-BE) driver to express *Ras*^*G12V*^ in the larval wing imaginal discs (Bharti et al., 2023; Singh et al., 2021), besides haltere and salivary glands **(Fig. S1A)**. As reported previously, the *Ras*^*G12V*^-expressing wing imaginal disc displayed excessive folding while maintaining intact actin cytoskeletal integrity, a hallmark of its hyperplastic growth (Karim and Rubin, 1998; Khan et al., 2013) (Actin, **Fig. 1A, A’**). This hyperplastic overgrowth is also evident in intact larvae **(Fig. 1B)** as it results in an increased overall disc size **(Fig. 1C)**. Examination of *Drosophila* E-cadherin (DE-cad, encoded in *shotgun*)—a component of adherens junctions (AJs) (Tepass et al., 1996)—and Fas III (Fasciclin-III)—a component of septate junctions (Wells et al., 2013)—further reveals an intact cellular polarity (**Fig. 1D-E**). Despite intact architecture, *Ras*^*G12V*^ discs displayed significantly elevated numbers of phospho-histone H3 (PH3)– positive cells (**Fig. 1F, G**). Together, these findings indicate hyperplastic overgrowth without overt loss of epithelial organization, consistent with a precancerous lesion. Given the emerging link between oncogenic signaling and metabolic reprogramming, we further assessed metabolic markers. For instance, *Ras*^*G12V*^ discs showed elevated eIF4E-binding protein (4E-BP), a downstream target of mTOR signaling (**Fig. S1B**) (Miron et al., 2001), and Acetyl-CoA Carboxylase (ACC), a key enzyme in the fatty acid synthesis pathway (Musselman and Kühnlein, 2018) (**Fig. S1C**), indicating a localized metabolic reprogramming.

**Fig. 1:**
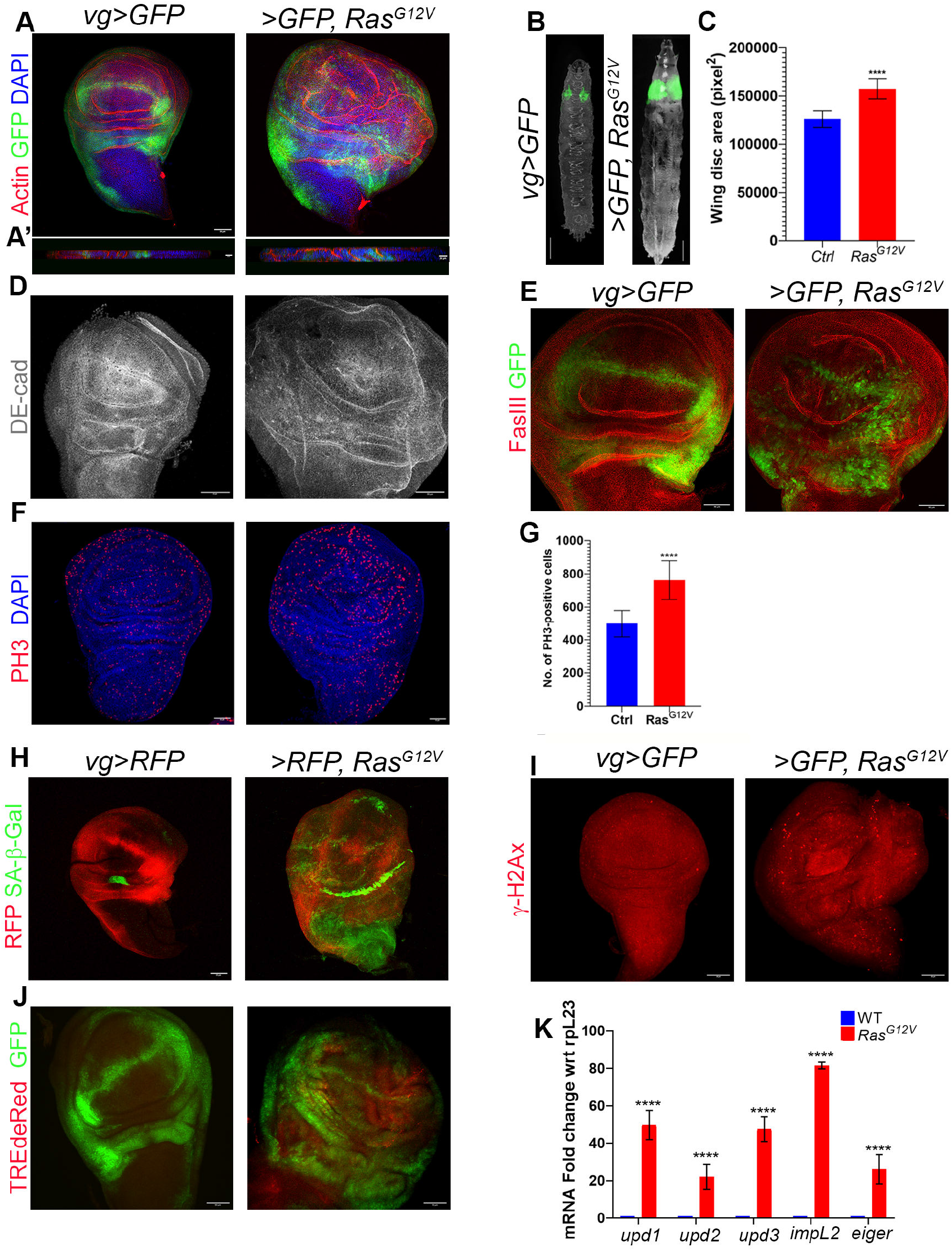
*Ras*^*G12V*^ expression in the wing imaginal disc triggers a precancerous lesion and OIS/SASP program. **A-A**^’^) Wing imaginal discs expressing *vg*>*GFP* (control) or *vg*>*GFP, Ras*^*G12V*^ displaying Actin (red) and DAPI (blue), and their Y-Z section (A^’^). GFP marks the *vg* domain. **B)** Representative larvae displaying GFP expression. **C)** Wing disc size quantification (N=10 wing discs, ^*\*\*\*\**^ *p<0*.*0001*, Data are presented as mean ± SEM). **D)** Wing disc displaying DE-Cad (grey) of the respective genotypes. **E)** Wing disc displaying Fas3 (red) of the respective genotypes. GFP marks the *vg* domain. **F-G)** PH3 (red, F) staining and its quantification (G). Nuclei are counterstained for DAPI (N=10 wing disc quantified, ^*\*\*\*\**^ *p<0*.*0001*, Data are presented as mean ± SEM). **H)** SA-β-Gal activity (green) in the respective genotypes. *vg* domain marked with RFP (*vg*>*RFP*). **I)** Wing disc displaying γ-H2Ax (red) of the respective genotypes. **J)** TRE-dsRed levels in the wing imaginal disc. *vg* domain marked with GFP. **K)** qPCR analysis of cytokine expression in the wing imaginal disc of respective genotypes (N=3 biological replicates, ^*\*\*\*\**^ *p<0*.*0001*, Data are presented as mean ± SEM). Scale bars: 50µm (A, D, E, F, H, I, and J); 100µm (B), and 20µm (A’).

We next investigated whether oncogenic stress triggers an OIS program. As shown previously (Ito and Igaki, 2016; Nakamura and Igaki, 2017), *Ras*^*G12V*^ discs exhibited robust SA-β-Gal activity **(Fig. 1H)—**a key senescence marker (Dimri et al., 1995)—and an elevated γH2Ax (**Fig. 1I**), indicative of DNA damage (Park et al., 2012). Further, increased TRE-dsRed reporter activity **(Fig. 1J)** reflects an activation of the JNK pathway (Chatterjee and Bohmann, 2012)—a key pathway associated with the induction of senescence and SASP (Garcia-Arias et al., 2023; Ito and Igaki, 2016). qPCR analysis further revealed altered expression of cytokines including *upd1*/*2/3, eiger*, and *impL2* **(Fig. 1K)**, consistent with induction of SASP. Importantly, *Ras*^*G12V*^ did not activate SA-β-Gal activity in the salivary gland, unlike the wing disc **(Fig. S1D)**, despite showing a *Ras*^*G12V*^-mediated organ hypertrophy. We noted increased expression of *upd2, upd3*, and *impL2* in the salivary glands **(Fig. S1E)**, demonstrating that *Ras*^*G12V*^ triggers a distinct cytokine profile independent of the OIS program.

Collectively, these data demonstrate that *Ras*^*G12V*^ induces a hyperplastic precancerous lesion that engages an OIS-mediated SASP, thereby promoting pro-inflammatory signaling.

### *Ras*^*G12V*^ activation in the wing disc induces systemic MetS in the host larva

Metabolic syndromes (MetS) are characterized by a cluster of disorders—namely, obesity, high blood pressure, hyperglycemia, and dyslipidemia—that are strongly associated with precancerous and early-stage cancers (Bashir et al., 2025; Belladelli et al., 2022; Esposito et al., 2013). We noted that *Ras*^*G12V*^-expressing larvae show increased body mass, both wet and dry weight **(Fig. 2A)**—reflecting true biomass accumulation, in contrast to the increase in wet weight commonly observed in the bloating phenotype of cancer-induced paraneoplastic nephrotic syndrome (Xu et al., 2024). Consistent with their biomass accumulation, *Ras*^*G12V*^-expressing larvae displayed an enlarged fat body **(Fig. 2B**)—the fly equivalent of mammalian adipose tissue and the liver (Musselman and Kühnlein, 2018)—revealing their obesity and elevated whole-body triacyl glyceride (TAG) **(Fig. 2C**), a marker for hyperlipidemia (Birse et al., 2010; Heier and Kühnlein, 2018; Musselman et al., 2013). However, we noted a decrease in circulating glucose and trehalose **(Fig. 2D)**, which is in contrast to the hyperglycemia observed in human MetS (Eckel et al., 2005). We further noted an elevated lipid storage marked by increased lipid droplet size (**Fig. 2E, F**), along with cellular hypertrophy **(Fig. 2G)**. This is further accompanied by elevated ACC (**Fig. 2H**). In addition, the *Drosophila* fat body also serves as the primary site for glycogen storage (Yamada et al., 2018). Consistently, we observed elevated glycogen accumulation as indicated by PAS staining **(Fig. 2I)**.

**Fig. 2:**
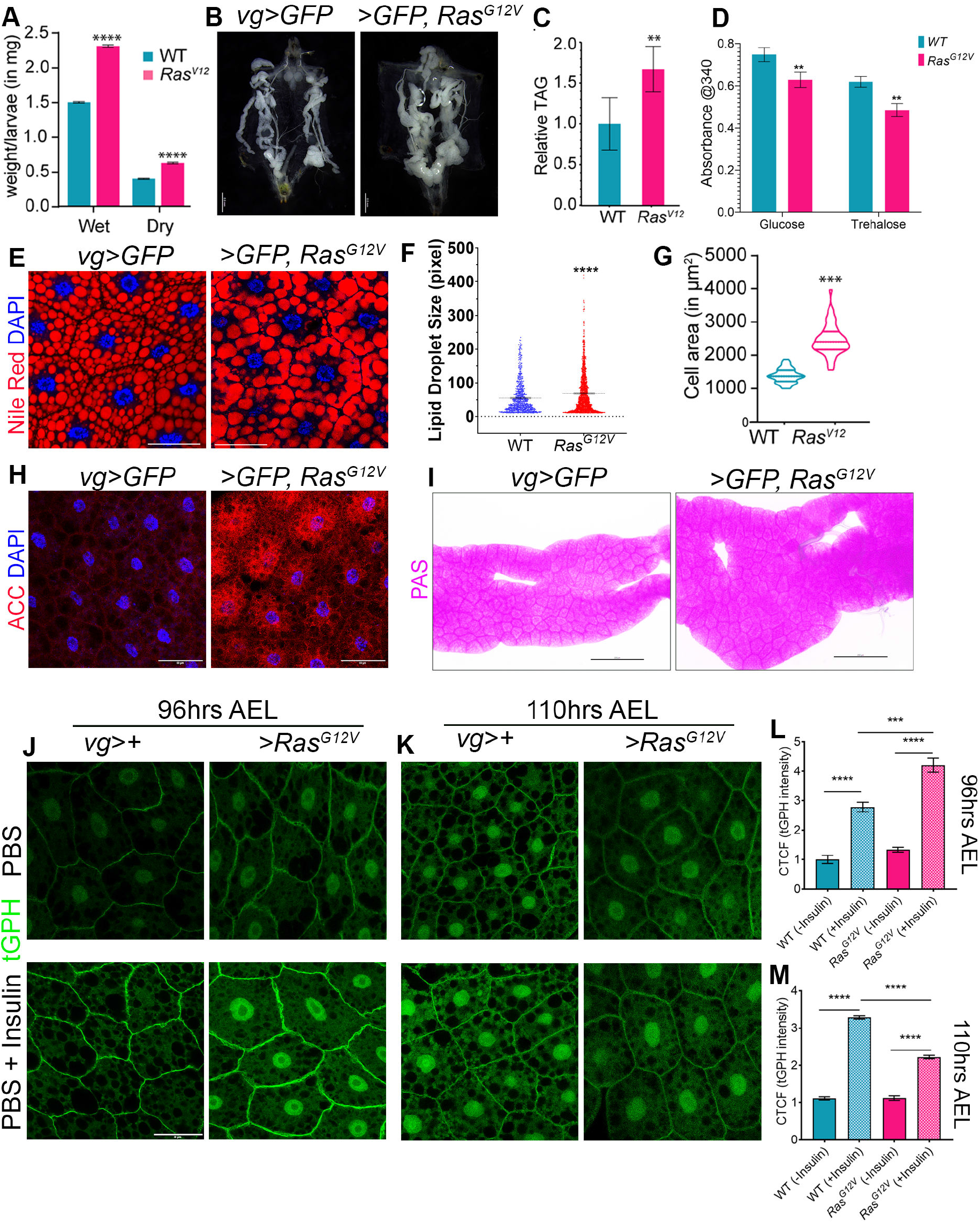
*Ras*^*G12V*^-expressing larvae display MetS phenotype. **A)** Wet and dry weight comparison of wild type and *Ras*^*G12V*^ expressing larvae (N=10 each replicate, 3 biological replicates for each genotype, **** *p<0*.*0001*, Data are presented as mean ± SEM). **B)** Fat body comparison of respective genotypes. **C-D)** Hemolymph TAG (C) and Glucose + Trehalose (D) levels from the respective genotypes. (N=10 each replicate, 3 biological replicates for each genotype, ** *p<0*.*01*, Data are presented as mean ± SEM). **E-F)** Nile red (red, E) for lipid droplet size in fat body and its quantification (F). Nuclei are stained for DAPI (blue) (N=10 fat body quantified, **** *p<0*.*0001*, Data are presented as mean ± SEM) **G)** Fat body cell size quantification (n=>100 cells quantified across 5 fat bodies per genotype; *** *p<0*.*001*, Data are presented as mean ± SEM). **H)** ACC (red) in the fat body of respective genotypes. Nuclei are stained with DAPI. **I)** PAS staining of the fat body of respective genotypes. **J-M)** tGPH levels in the fat body upon incubation with PBS (upper panel) or PBS + Insulin (lower panel) (96hrs AEL (J) and its quantification (L) and 110hrs AEL (K) and its quantification (M)). (N=10 larvae per genotype per time point, **** *p<0*.*0001* and ** *p<0*.01, Data are presented as mean ± SEM). AEL: After Egg Laying. Scale bar: 0.5mm (B), 50µm (E, H, J, and K), and 200µm (I).

Obesity and associated chronic inflammation induce insulin resistance (McArdle et al., 2013; Qatanani and Lazar, 2007). In *Drosophila*, active insulin signaling is assayed using a PI3K activity reporter, in which PIP3 induces membrane recruitment of a GFP-tagged Pleckstrin homology domain (tGPH) (Britton et al., 2002). In the presence of insulin, insulin-sensitive cells display membrane localization of tGPH, while insulin-resistant cells do not. Thus, the fat body from larvae starved for 6 hr from both genotypes, wild-type and *Ras*^*G12V*^-expressing larvae (mid-third instar, 96 ± 2 hrs after egg laying (AEL)) (with no ongoing insulin signaling), displays no tGPH membrane-localization **(Fig. 2J, L)**. Upon incubation of fat body with human insulin (Musselman et al., 2011; Pasco and Léopold, 2012), *Ras*^*G12V*^-expressing larval fat body displayed increased membrane tGPH localization as compared to wild type **(Fig. 2J, L)**, suggesting an elevated insulin signaling and higher responsiveness in the fat body. In contrast, in the late third instar stage (110 ± 2hrs AEL), wild-type fat bodies remained responsive to insulin signaling, showing robust tGPH enrichment upon stimulation (**Fig. 2K, M**), whereas *Ras*^*G12V*^-expressing larvae displayed a lower response to the exogenous insulin signaling—a phenomenon seen during insulin resistance conditions. These observations, together, suggest a hyperinsulinemia-to-insulin resistance transition observed upon *Ras*^*G12V*^ expression.

The majority of these *Ras*^*G12V*^-expressing larvae die during the larva-to-adult transition, with ~5% larvae transitioning into the adult **(Fig. S2A)** and reduced adult survival, with the majority dying within 8 days post-eclosion **(Fig. S2B)**. Eclosed adult display deformed wing **(Fig. S2C)**, loss of fat body (Nile red, second panel, **Fig. S2C)**, loss of muscle mass (Actin, third panel, **Fig. S2C**), along with organ atrophy, as suggested by ovary degeneration (fourth panel, **Fig. S2C)**. These results, together, demonstrate a MetS-cachexia transition in *Ras*^*G12V*^-expressing adults.

These results reveal that a precancerous epithelium displaying *Ras*^*G12V*^ signaling systemically perturbs energy homeostasis in the host animal, marked by a MetS-like phenotype—obesity, hyperlipidemia, and a hyperinsulinemia-to-insulin resistance transition (Alberti et al., 2005; Eckel et al., 2005; Huang, 2009). Interestingly, we noted a MetS-to-cachexia transition, further confirming MetS as a precursor to cachexia (Wyart et al., 2025).

### OIS program in the wing disc activates a senescence-like program in the fat body via Upd1

An interesting aspect of senescence/OIS induction is its paracrine senescence: a senescence-induced senescence phenomenon (Acosta et al., 2013; Admasu et al., 2021; Kuilman et al., 2008; Nelson et al., 2012), mainly mediated by the inflammatory cytokines. However, SASP factors can travel through the bloodstream and may act on distant organs (Schafer et al., 2020). A recent study by Kiourtis et al., 2024 confirms these assumptions, showing that senescent hepatocytes in the mouse liver can activate a senescence relay to distant organs, via TGF-β, leading to multi-organ dysfunction (Kiourtis et al., 2024). We tested this observation and noted an elevated SA-β-Gal activity in the fat body of *Ras*^*G12V*^-expressing larvae **(Fig. 3A)**. Further, we noted an increased JNK signaling in the fat body (TRE-dsRed, **Fig. 3B**) and an elevated ROS activity (DHE, **Fig. 3C**), confirming a senescence-like state in the fat body. This further led to increased expression of *upd2* and *upd3* from the fat body **(Fig. 3D)**.

**Fig. 3:**
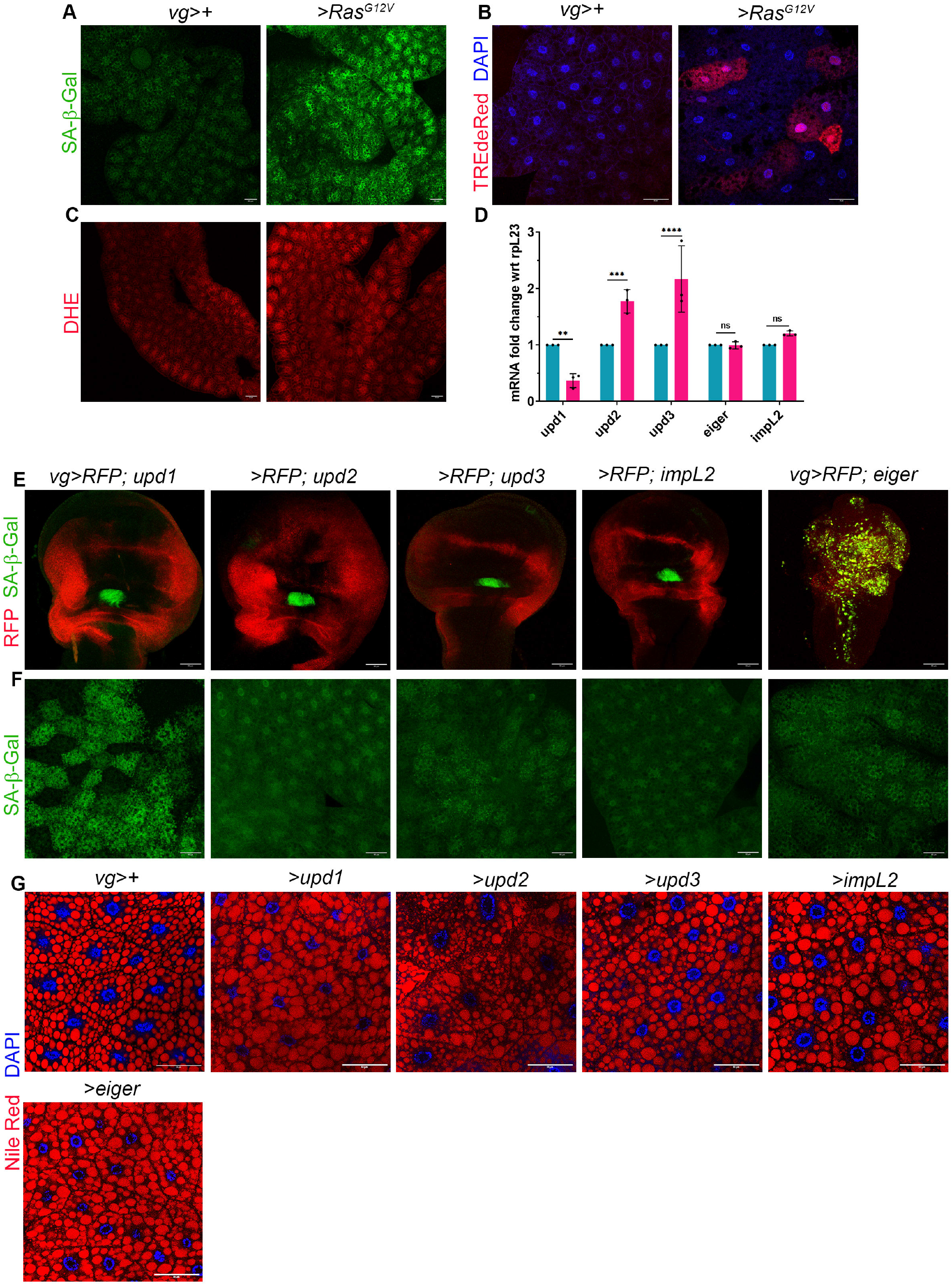
Larval fat body displays activation of Upd1-mediated senescence-like program. **A-C)** SA-β-Gal activity (A), TRE-dsRed levels (B), and DHE levels (C) in wild type and *Ras*^*G12V*^-expressing larval fat body (N=10 per genotype). **D)** qPCR analysis of cytokine levels in the fat body of the respective genotypes. (\*\*\*\**;p<0*.*0001*, *** *p<0*.*001*, and ** *p<0*.*01*, Data are presented as mean ± SEM). **E-F)** SA-β-Gal activity (green) in the wing disc (E) and the fat body (F) upon ectopic expression of different cytokines from the wing imaginal disc. RFP marks the *vg*-domain in the wing imaginal disc. **G)** Nile red staining in fat body upon *upd1/2/3, impL2*, and *eiger* expression from the wing imaginal disc. Scale bar: 50µm.

To identify key senescence-relay cytokines, we overexpressed individual SASP candidates in wing imaginal discs using the *vg*-*Gal4* driver and assessed fat body senescence.

Overexpression of *upd1* induced robust ectopic senescence in the distant larval fat body **(Fig. 3E, F)**. Notably, *eiger* triggered autonomous SA-β-Gal activity in the wing imaginal disc **(Fig. 3E)**. Consistently, ectopic expression of *upd1* from the adult male accessory gland (MAG) also induced senescence in the adult fat body **(Fig. S3A, B)**. These data position *upd1* as the primary relay cytokine linking *Ras*^*G12V*^-mediated OIS program to the distant fat body. Interestingly, *upd1* overexpression also promotes lipid accumulation in the fat body **(Fig. 3G)**, phenocopying the lipid accumulation observed upon *Ras*^*G12V*^ expression. Together, these observations suggest that *Ras*^*G12V*^-driven OIS initiates a distant senescence relay through SASP cytokines—primarily *upd1*—that propagates senescence and metabolic reprogramming to the distant fat body. However, we did not observe any increase in the JAK-STAT signaling levels in the wing disc or the fat body upon *Ras*^*G12V*^ expression (**Fig. S3C**), suggesting that the activation and relay of senescence do not involve JAK-STAT signaling.

Senescence induction and its associated pathophysiology are context-dependent phenomena, as not all oncogenes trigger OIS in every context (Kwiatkowska et al., 2023). To assess context dependency of OIS and its relay in *Drosophila*, we expressed distinct oncogenes (*Ras*^*G12V*^, *Yki*^*3SA*^, and *N*^*intra*^) in three different cell types, including proliferating and specified wing disc cells (using *vg*-*Gal4*), determined and non-dividing eye disc cells (using *GMR*-*Gal4* (Li et al., 2012)), and postmitotic adult male accessory gland (MAG) primary cells (using *ov*-*Gal4* (Bhattacharya et al., 2024; Kumari and Sinha, 2021)). As shown previously, *Ras*^*G12V*^ induced senescence only in proliferating wing disc cells (see **Fig. 1H**) but not in non-dividing eye disc or postmitotic MAG cells **(Fig. S4B, C)**. In contrast, *Yki*^*3SA*^ triggered senescence exclusively in MAG cells **(Fig. S4A-C)**, whereas *N*^*intra*^ failed to induce senescence in all contexts tested **(Fig. S4A-C)**. Notably, the neoplastic model *scrib*-*IR*; *Ras*^*G12V*^ induced senescence across all cellular contexts (**Fig. S4A–C**). These findings indicate that oncogene-induced senescence is highly context-dependent, whereas neoplastic transformation overrides cell-state restrictions.

We further observed that senescence relay is also context-dependent. Senescence originating from wing and eye imaginal discs propagated to the fat body, whereas MAG-derived senescence did not (**Fig. S4A–C**). Moreover, relay efficiency also depended on the secondary organ, as wing disc–derived senescence relay failed to propagate to larval muscle (**Fig. S4D**).

Together, these results demonstrate that *Ras*^*G12V*^-driven OIS in the wing disc induces a context-dependent Upd1-mediated systemic senescence relay that propagates senescence and metabolic alterations to the distant fat body, independently of canonical JAK–STAT activation.

### Systemic *Ras*^*G12V*^ activation triggers IPCs, NPF, and Hugin neuron activity

Hyperphagia or prolonged caloric intake is known to induce MetS-associated weight gain, obesity, insulin resistance, and hyperglycemia. Hence, we analyzed the feeding profile of *Ras*^*G12V*^-expressing larvae by ingestion of Carmoisine red mixed colored food (Agrawal et al., 2009), and found an increased food intake **(Fig 4A, B)**, confirming hyperphagia. In *Drosophila*, the NPF and the Hugin neurons play pivotal roles in regulating feeding behavior (Nässel and Zandawala, 2020). When checked for NPF—that promotes feeding—and Hugin—that suppresses feeding—expression in the brain, we found an increased NPF and decreased *hugin* expression in *Ras*^*G12V*^-expressing larvae **(Fig 4C)**. In addition, we also noted an increased *dILP2, 3*, and *5* expression in *Ras*^*G12V*^-expressing larval brains **(Fig. 4D)**. This is further confirmed by elevated dILP2 reporter expression in insulin-producing cells (IPCs) neurons in the brain (*dILP2mCherry*, **Fig. 4E)**.

**Fig. 4:**
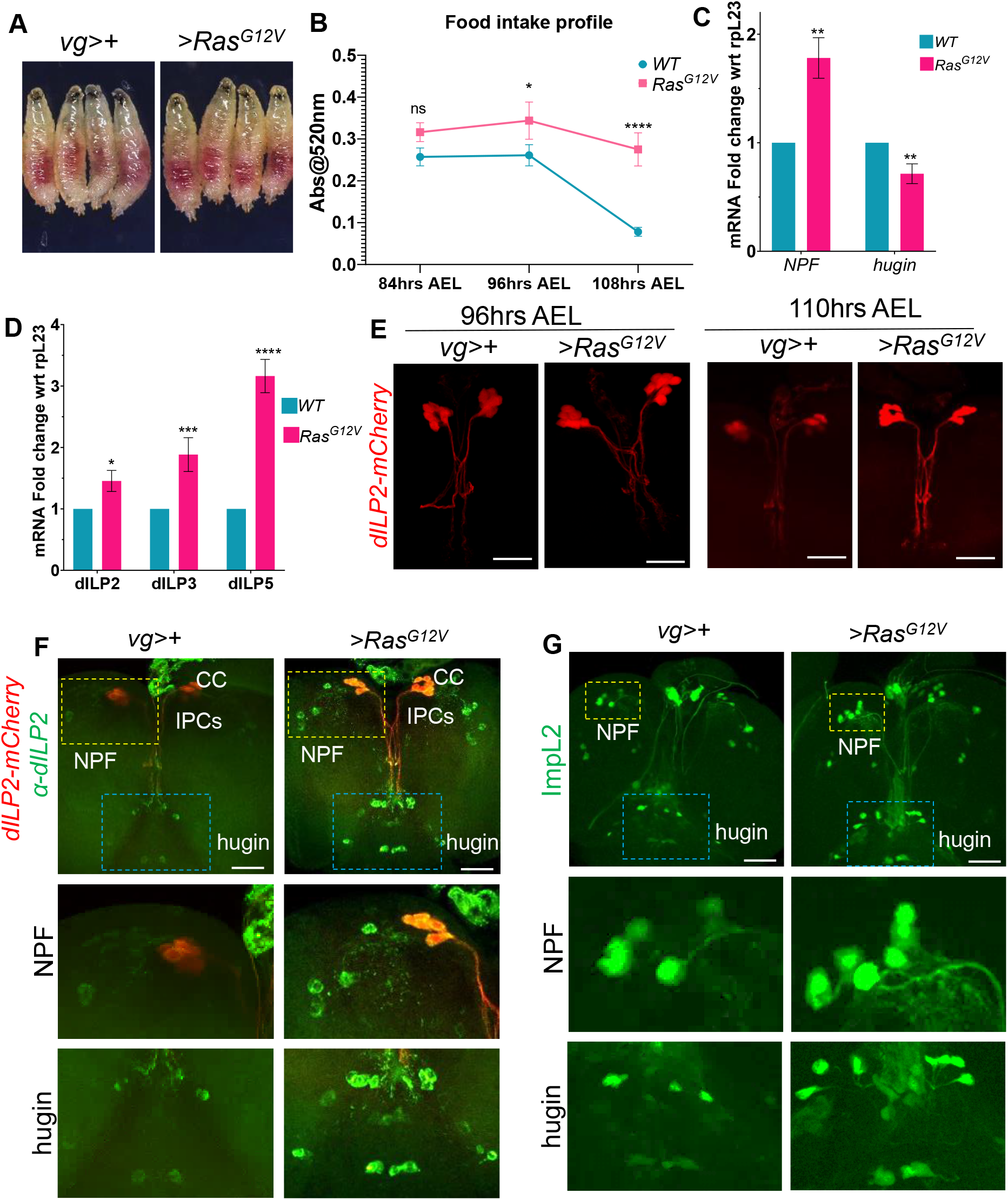
Epithelial *Ras*^*G12V*^ communicates with IPC neurons to promote hyperphagia and hyperinsulinemia. **A)** Brightfield images of wild-type and *Ras*^*G12V*^-expressing larvae after feeding on carmoisine red-colored food. **B)** Quantification of food intake by measuring the absorbance of carmoisine dye from larval homogenates at 520/280 nm. (N=10 each replicate, 3 biological replicates for each genotype, **** *p<0*.*0001*, and \**;p<0*.1, Data are presented as mean ± SEM). **C)** qPCR to quantify expression of sNPF, and hugin, from the brain of respective genotypes (*\*\*p<0*.*01*, Data are presented as mean ± SEM). **D)** qPCR analysis in the brain samples of respective genotypes (**** *p<0*.*0001*, *** *p<0*.*001*, and * *p<0*.1; Data are presented as mean ± SEM). **E)** Brain expression of dILP2-mCherry reporter of 96hrs and 110hrs AEL larvae of respective genotypes. (N=10 larval brains analyzed for each genotype). **F)** Expression of dILP2 in sNPF and hugin neurons, besides corpora cardiaca of respective genotypes. (N=10 larval brains analyzed for each genotype). **G)** Expression of ImpL2-GFP in sNPF and hugin neurons of respective genotypes. (N=10 larval brains analyzed for each genotype). Scale bars: 1mm (A), 50µm (D, F, and G).

Previous studies have shown an intricate involvement of NPF and Hugin neurons with IPCs through paracrine insulin signaling to regulate feeding (Kapan et al., 2012; Lee et al., 2004; Melcher et al., 2006) and release of insulin from insulin-producing cells (Nässel et al., 2013; Rajan and Perrimon, 2012; Sudhakar et al., 2020). We observed increased accumulation of dILP2 in NPF and Hugin neurons (Fig. 4F), suggesting crosstalk between them. These neurons respond to insulin by expressing the IGF-binding protein, ImpL2, to activate downstream insulin signaling (Bader et al., 2007; Bader et al., 2013; Melcher et al., 2006); therefore, *impL2*-GFP expression is a proxy for the activity of these neurons. We found upregulation of *impL2*-*GFP* in NPF and Hugin neurons (**Fig. 4G**), which is responsible for increased dILP2 uptake by these neurons **(Fig. 4F)**.

In *Drosophila*, GABAergic neurons regulate dILP secretion by the IPCs (Rajan and Perrimon, 2012). The upregulation of JAK-STAT signaling in these GABAergic neurons alleviates the inhibitory restraint on the secretion of dILPs from IPCs (Rajan and Perrimon, 2012). As expected, we found elevated STAT levels in the brains of *Ras*^*G12V*^-expressing larvae compared to controls **(Fig. S5A)**.

### Senescence induction in the fat body precedes MetS-associated lipid accumulation, and its suppression mitigates the MetS phenotype in the fat body

Stage-wise Nile red and SA-β-Gal activity in the fat body reveals an early activation of the senescence program in the fat body **(Fig. 5A, B)** of the *Ras*^*G12V*^ expressing larvae, even before the onset of MetS-associated lipid accumulation **(Fig. 5A, B)**. In contrast, in the neoplastic tumor model of *scrib-IR; Ras*^*G12V*^ (Brumby and Richardson, 2003) larval fat body displays activation of the senescence program and increased lipid accumulation droplet size during both early and late stages **(Fig. 5A, B)**, which is consistent with the early onset of cachexia phenotype (Figueroa-Clarevega and Bilder, 2015; Hodgson et al., 2021; Kwon et al., 2015; Lodge et al., 2021). Thus, it is likely that, in *Ras*^*G12V*^-expressing larvae, senescence induction in the fat body precedes the MetS phenotype.

**Fig. 5:**
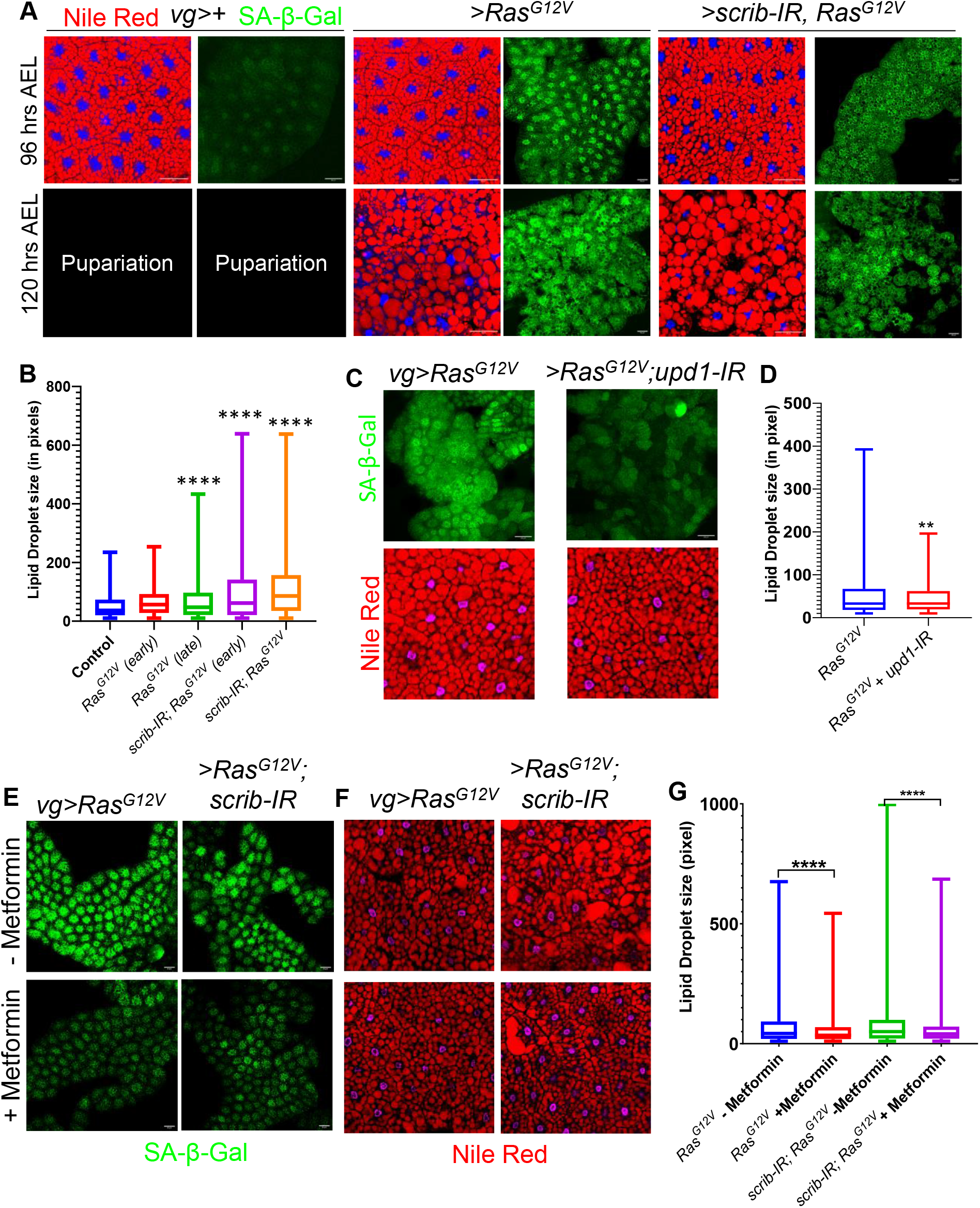
Senescence precedes MetS, and its suppression reduces MetS and cachexia. **A)** Nile red (red) and SA-β-Gal activity (green) in the fat body in precancerous *Ras*^*G12V*^ and neoplastic *scrib-IR*; *Ras*^*G12V*^ models at 96hrs and 120hrs AEL. **B)** Lipid droplet size quantification of respective genotypes (N=10 fat body analyzed, **** *p<0*.*0001*, Data are presented as mean ± SEM). **C)** SA-β-Gal activity (green, upper panel) and Nile red (red, bottom panel) in the larval fat body upon genetic knockdown of *upd1* in the *Ras*^*G12V*^-expressing larvae (*vg Gal4*>*UAS-Ras*^*G12V*^, *UAS-upd1-IR*) as compared to *Ras*^*G12V*^-expressing larvae alone (*vg Gal4>UAS-Ras*^*G12V*^). **D)** Lipid droplet size quantification for (C) (N=10 fat body analyzed per genotype ** *p<0*.*01*, Data are presented as mean ± SEM). **E-F)** SA-β-Gal activity (green, E) and Nile red (red, F) in the fat body with metformin (lower panel) and without metformin (upper panel) in precancerous *Ras*^*G12V*^ and neoplastic *scrib-IR*; *Ras*^*G12V*^ models. **G)** Quantification of lipid droplet size from (F) (N=10 fat body analyzed, \*\*\*\**p<0*.*0001*, Data are presented as mean ± SEM). Scale bar: 50µm.

Interestingly, knockdown of *upd1* in the *Ras*^*G12V*^-expressing wing disc reduced SA-β-Gal activity **(Fig. 5C)** and restored lipid droplet size **(Fig. 5C, D)**, demonstrating that disruption of senescence relay to the fat body mitigates the MetS-associated phenotype and further increases host survival **(Fig. S6A)**.

To further understand the mitigation of senescence relay and its impact on MetS phenotype, we employed a pharmacological approach. Metformin, a biguanide derivative, is widely used to treat type 2 diabetes (for review, see (Lv and Guo, 2020; Nasri and Rafieian-Kopaei, 2014)) and has increasingly been recognized for its anticancer and anti-aging benefits (for review, see (Zhu et al., 2024)). Metformin acts as a senomorphic agent—modulating the SASP and restraining inflammatory signaling without eliminating senescent cells (Abdelgawad et al., 2023). Interestingly, feeding Metformin to the larvae bearing *Ras*^*G12V*^ or *scrib-IR; Ras*^*G12V*^-expressing larvae, lowered SA-β-Gal activity in their fat body **(Fig. 5E)** along with a decrease in lipid droplet size **(Fig. 5F, G)**. Interestingly, in *scrib-IR; Ras*^*G12V*^ larvae, Metformin also reduced tumor size **(Fig. S6B, C)**, without altering the tumor’s senescence status **(Fig. S6B)**.

Together, these findings demonstrate that Upd1-dependent senescence relay triggers early fat-body senescence and MetS, establishing a systemic axis that promotes MetS progression toward cachexia and can be therapeutically modulated by senomorphic intervention.

## DISCUSSION

The relationship between metabolic syndrome (MetS) and cancer has long been recognized in clinical and epidemiological studies, yet the direction of causality remains unresolved (Bashir et al., 2025; Belladelli et al., 2022; Esposito et al., 2013; He et al., 2016; Zhang et al., 2023b). MetS has traditionally been considered either a risk factor for tumorigenesis or a metabolic consequence of established malignancies (Belladelli et al., 2022). Our findings provide experimental evidence that precancerous oncogenic lesions, by activating the OIS/SASP program, can initiate systemic MetS. Using a *Drosophila* model of epithelial *Ras*^*G12V*^ activation, we demonstrate that local OIS in a hyperplastic precancerous lesion can propagate senescence to the distant fat body and reprogram systemic metabolism.

A key insight from this study is that senescence signaling is not restricted to the local level alone (Acosta et al., 2013; Nelson et al., 2012), but can also be relayed to distant tissues. Although previously speculated, the mechanistic underpinning of this phenomenon has recently been explored in a mouse model, which revealed that senescence in hepatocytes can relay senescence to distant organs via TGF-β signaling (Kiourtis et al., 2024). Our study demonstrates that *Ras*^*G12V*^-induced senescence in the wing imaginal disc triggers a SASP program that includes the IL-6–like cytokine Upd1, which relays senescence signals to the distant fat body. Our findings extend this concept by demonstrating that both the induction of OIS and its systemic relay are highly context-dependent processes: not all oncogenes can trigger senescence, not all senescence events propagate systemically, and not all recipient tissues are permissive to secondary senescence. Such selectivity may help explain why metabolic disturbances arise in specific organs in an oncogene-specific manner.

Importantly, relay of senescence to the fat body precedes its overt manifestation of MetS, suggesting that senescence drives metabolic reprogramming rather than the converse. Thus, the fat body exhibited early senescence markers, followed by lipid accumulation and hypertrophy, characteristic signatures of its onset of MetS (Alberti et al., 2005). In parallel, *Ras*^*G12V*^-expressing animals displayed hyperphagia and increased production of insulin-like peptides (dILP2/3/5), indicating a chronic state of hyperinsulinemia (Belladelli et al., 2022) that ultimately progressed to insulin resistance (Roberts et al., 2013). These findings suggest that SASP-associated Upd1 may reshape systemic energy homeostasis by influencing both peripheral metabolic tissues and central feeding circuits.

Finally, we demonstrate that blocking OIS relay can mitigate systemic MetS. Thus, genetic knockdown of *upd1* in *Ras*^*G12V*^-expressing epithelium significantly reduced fat-body senescence and lipid accumulation, while pharmacological intervention with the senomorphic drug Metformin attenuated both senescence and metabolic defects. These findings support a model in which OIS-driven SASP signaling links early oncogenic lesions to host metabolic dysfunction, ultimately predisposing the organism to cachexia-like wasting **(Fig. 6)**. Importantly, this axis appears to be therapeutically tractable.

**Fig. 6:**
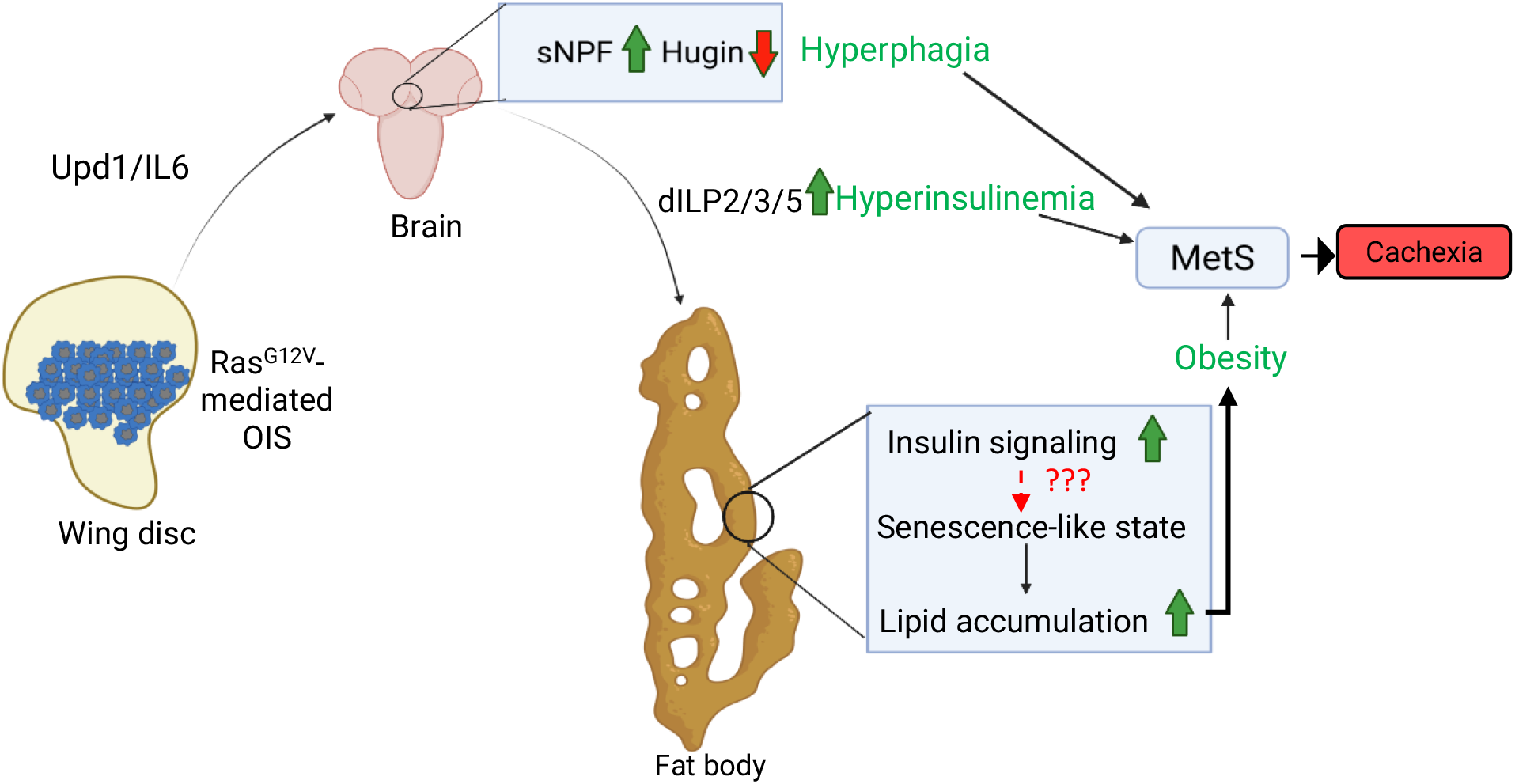
OIS-MetS axis in *Ras*^*G12V*^-precancerous lesion leads to cachexia.

Together, our work reveals that the OIS program in precancerous lesions can act as an early systemic signaling hub that reshapes host metabolism, establishing a mechanistic bridge between oncogenic stress and MetS. These findings suggest that metabolic dysfunction observed in cancer patients may, in part, originate from early oncogenic lesions long before tumor detection. Targeting the senescence–metabolic axis may therefore represent a promising strategy to restrain tumor progression on the one hand, and its detection may serve as an early diagnostic marker for cancers, such as CRC, on the other.

## MATERIALS AND METHODS

Fly lines were cultured on standard yeast-cornmeal agar at 25°C and 29°C for RNAi lines. Synchronized egg-laying was performed at 25°C for 4h to collect larvae at the same developmental stage. Larval tissues were dissected in cold PBS, fixed in 4% paraformaldehyde, stained with the desired antibody, and mounted. Image acquisition was done in various confocal microscopes, and data analysis was performed using FIJI. Detailed staining protocol, quantification methods, and statistical analysis are provided in the **Supplementary Information**. All the genetic stocks and reagents used have been procured from public repositories and as gifts from other researchers (**Supplementary Information, SI Table 1 and 2**).

## ACKNOWLEDGMENTS

IIT Kanpur supports research in the PS laboratory. We thank UGC, CSIR, and MHRD for financial support to JT, SSP, SK, SB, and IMA.

## AUTHOR CONTRIBUTIONS

JT, SSP, and PS designed the research; JT and SSP performed the research, and SB and IMA contributed; SK and JT conducted molecular tests. SSP and PS wrote the manuscript, and all authors reviewed it.

