## Supplementary Information for "*Ras^G12V^* Oncogene-Induced Epithelial Senescence and Its Relay Promotes Host Metabolic Syndrome in *Drosophila*"

This PDF file includes:

A. Supplementary Text
B. Supplementary Table (S1-S4)
C. Supplementary References
D. Supplementary Figure legends (Fig. S1-S6)
E. Supplementary Figures (Fig. S1-S6)

**SUPPLEMENTARY TEXT**

**EXPERIMENTAL METHODS**

***Drosophila* Culture and Genetic Crosses.** For expression in wing, boundary enhancer-driven *vestigial* driver (*vg-Gal4)* (Bharti et al., 2023; Williams et al., 1991) was used, *GMR-Gal4* (Li et al., 2012) for eye expression and *ov-Gal4* (Bhattacharya et al., 2024; Heifetz et al., 2000) for expression in MAG. (see **SI** **Table 1 and 2** for detailed stocks and resources). *Drosophila* cultures were reared on standard cornmeal agar at 25 ± 0.5 °C. Synchronised 4 h egg lays were used to collect stage-matched larvae; 50 larvae per genotype were cultured in six replicate vials. For adult experiments, same-day eclosed flies (~30 per genotype) were collected and maintained. Flies were transferred to fresh food vials every 3–4 days to prevent mortality due to entrapment in liquefied food. For RNAi-based studies, first instar larvae or newly eclosed adults of the desired genotypes were maintained at 29 ± 0.5 °C (Duffy, 2002).

**Immunohistochemistry and Imaging.** Mid-third instar (96 ± 2 hrs after egg laying (AEL)) and late-third instar larvae (110 ± 2hrs AEL) were fixed in 4% PFA in PBST (PBS + 0.2% Triton X-100) for 40 minutes. For adult male accessory glands (MAGs), anaesthetised adult males were dissected in cold PBS, and the MAG was gently pulled from the abdomen and fixed. Samples were then washed (PBST x3), blocked (10% BSA, 2h) and incubated with primary antibody (see **SI Table 1**) overnight at 4°C. Samples were again washed (PBST x3) and incubated with secondary antibody for 4h at RT in dark, washed (PBS x3), counterstained with Actin (Phalloidin) or a nuclear stain (DAPI) for 1h, washed (PBS x3), and mounted in Vectashield. For staining of reporter lines, tissues were fixed (4% PFA, 40min), counterstained (DAPI) and mounted for imaging.

For whole larval fat body and muscle dissection**,** third instar larvae were immobilised dorsally on a wax plate using tungsten needles as described previously (Hodgson et al., 2021; Lodge et al., 2021). A longitudinal incision along the anterior-posterior axis was made using micro-scissors to expose larval fat body and muscle. The overlaying cuticle was spread and pinned to form a flat sheet. Tissues were fixed in 4% PF/PBS (30mins), followed by three PBS washes. The cuticle with the fat body was transferred to slides for imaging. For muscle visualisation, the attached fat body and other tissues were removed, and muscles were stained with Phalloidin for 2 hours, washed (PBS x3) and mounted in antifade glycerol medium.

*Drosophila* larvae were rinsed in PBS and fixed overnight in 4% PF in PBST. Adult flies were imaged under anaesthesia. Brightfield imaging of larvae, pupae, and adults was done using a Leica M205 FA stereomicroscope. Confocal images were acquired with a Leica SP6, Leica Stellaris, Zeiss LSM700, or AXR Nikon. Images were analyzed and processed using Leica confocal software, LAS AF, or FIJI.

**Quantification and Statistics.** All image acquisitions used for tGPH fluorescence intensity measurements were performed using identical settings for both control and experimental samples. Fluorescence intensities were quantified in FIJI. The freehand selection tool was used to mark the ROI for calculating integrated densities and surface area within the cell boundaries and the background. Corrected total cell fluorescence (CTCF) was calculated as: CTCF=Integrated density − (Cell area × Mean background fluorescence).
For cell area quantification, the freehand tool was used to mark the cell boundaries to calculate the surface area.

Quantitative data were plotted, and statistical analysis was performed using GraphPad Prism 8.0 to calculate p-values. Data are expressed as mean ± standard error of the mean (SEM). Comparisons between two groups were made using a two-tailed unpaired t-test. For multiple group comparisons, one-way ANOVA and two-way ANOVA, followed by post hoc multiple comparison tests, were employed. Kaplan-Meier survival curves were plotted using cumulative mortality data. Survival curves were analysed using the log-rank (Mantel-Cox) test. A cutoff of p<0.05 was used to define statistical significance in all graphical plots. All images were assembled using Adobe Photoshop CS-6. Illustrations were made using Bio Render.

**Nile red staining and droplet quantification.** Larval fat bodies were dissected in chilled PBS, fixed in cold 4% PF in PBS (20 minutes), washed (PBS x3) and incubated in 0.005% Nile Red (1:200 dilution in PBS) for 1h at RT. Samples were washed (PBS x3) and incubated with TO-PRO-3 (1:1000 in PBS) for 1 hr at RT. Stained tissues were mounted in an antifade mounting medium and imaged using a confocal microscope (Musselman et al., 2011).
For lipid quantification the maximum-intensity projected images were converted to greyscale with consistent brightness and contrast and background subtraction (rolling ball radius ~50px). A consistent threshold was used, followed by watershed-based binary segmentation to separate clumped lipid droplets. To analyse the particles, the following parameters were set: Size: 10-infinity μm^2^; Circularity: 0.5-1.00. The data are then plotted on GraphPad Prism.

**Larval wet and dry weight estimation.** Late-third instar larvae (110 ± 2hrs AEL) were grouped (10 larvae per group; total 5 groups), rinsed with distilled water, gently blotted on Kimwipes, and transferred into pre-weighed 1.5 mL tubes to determine the mean wet weight per larva. Further, for dry weight, these larvae or adults were dried for 24 hrs at 60°C and reweighed. Wet weight represents the total biomass, including water content; dry weight reflects the biomass excluding water; the difference corresponds to water loss, indicating total body water content (Testa et al., 2013).

**Periodic acid–Schiff (PAS) staining.** Larval fat body were dissected (1% BSA in PBS), fixed in 4% PF/PBS for 20 minutes at RT, washed (1% BSA in PBS x2), treated with periodic acid for 5 minutes, and washed again. Subsequently, samples were incubated with Schiff’s reagent for 15 minutes, washed (1% BSA/PBS x2), and mounted in 80% glycerol diluted in PBS and imaged (Onkar et al., 2023).

**Dihydroethidium (DHE) staining**. *Drosophila* third instar larvae were dissected in cold Schneider’s Medium to isolate the fat body tissue. The dissected tissues were immediately incubated in Schneider’s medium containing 30 μM DHE for 5–7 minutes (Owusu-Ansah et al., 2008) at RT in the dark. Tissues were rinsed twice with PBS, mounted in antifade medium and imaged immediately.

**SA-β-Gal activity detection in larval and adult tissues.** Two methods for detection of SA-β-Gal activity were implemented. For the histochemical-based detection, the standard protocol was followed as described (Nakamura and Igaki, 2017). In brief, tissues were dissected (PBS), incubated in the 1X Fixative buffer (15mins, RT), washed (PBS x3), incubated with Staining Mixture, for 90 hrs in a 37^o^C water bath, in dark, washed (PBS x3) and mounted in 80% glycerol and imaged.
Fluorescein-based detection of SA-β-Gal activity was done as previously described (Garcia-Arias et al., 2023). In brief, tissues were dissected (PBS), fixed in 4% PF (30 minutes at RT), washed (1% BSA in PBS x 3), and incubated with the green probe (1:1000 dilution) for 2h at 37^o^C. Samples were washed (PBS x3) and mounted using antifade media and imaged immediately.

**Real-Time Quantitative PCR (RT-qPCR).** Total RNA was extracted using the TRIzol RNA extraction method (Rio et al., 2010) and quantified. 1μg of total RNA from each sample was reverse transcribed into cDNA. qPCR was conducted using PowerUp SYBR Green Master on a QuantStudio™ 5 Real-Time PCR System. Each reaction included 50ng of cDNA and gene-specific primers. Thermal cycling conditions were 95°C for 10 min, followed by 40 cycles of 95 °C for 15 s and 60 °C for 1 min. RPL23 was used as a housekeeping gene. Relative expression was calculated using the 2–ΔΔCt method from three biological replicates per genotype. Primers used in qPCR are detailed in **SI Table 3**.

**Triacylglycerol (TAG) quantification.** Groups of 10 age-matched third instar larvae (in 3 technical replicates for each of the two biological replicates) were homogenized in 100μl of PBST (PBS + 0.05% Tween-20) using a homogenizer. Homogenates were heat-inactivated at 70°C for 10 min, cooled, and centrifuged at 13,000xg for 3 min. Supernatants (20μl) were incubated with 200μl of Triglyceride Reagent at 37°C for 30 min. Glycerol released was measured using the Free Glycerol Reagent, and absorbance was read at 540 nm (SpectraMax® Mini Multi-Mode Microplate Reader, Molecular Devices). TAG concentration was calculated by subtracting free glycerol values from total glycerol values, normalized to protein content (Tennessen et al., 2014).

**Trehalose and Glucose quantification.** Mid-third instar larvae (*n=*10) were homogenized in 100μl cold Trehalase Buffer, TB (5 mM Tris pH 6.6, 137 mM NaCl, 2.7 mM KCl). Homogenates were heat-inactivated at 70 °C for 10 min, centrifuged, and supernatants collected. Samples (30μl) were incubated with either TB or Trehalase enzyme stock (3μl porcine trehalase per ml TB) at 37 °C for 24 h. Glucose was measured using a hexokinase-based assay, and absorbance was read at 340 nm (SpectraMax® Mini Multi-Mode Microplate Reader, Molecular Devices). Trehalose content was calculated by subtracting background glucose from trehalase-digested values (Tennessen et al., 2014)

**Quantification of larval food intake.** Larval feeding was quantified using a dye-based ingestion assay (Agrawal et al., 2009). A yeast paste containing Carmoisine Red dye (2 mg dye per gram of yeast) was placed centrally on 60 mm Petri dishes plated with 2% agar (prepared in distilled water). Synchronized larvae at defined time points (i.e., 84, 96, and 108hrs AEL) were transferred in groups of 25 onto the dyed paste and allowed to feed for 2 hours at 25 +0.5^o^C. Post-feeding, larvae were rinsed with distilled water, blotted dry, and snap-frozen in liquid nitrogen. Samples were homogenized in 500μL chilled PBS, kept on ice, and centrifuged at 15,000 rpm for 10 minutes at 4°C. The absorbance of supernatant was measured at 520 nm (SynergyTM H4, BioTek® Instruments, USA) to quantify dye ingestion. Absorbance at 280 nm was used for protein, and normalized food intake was displayed as Abs520/Abs280, *n=*4.

**Feeding larvae on metformin.** Synchronized *Drosophila* larvae were transferred at the second instar stage (∼48 hrs AEL) to standard cornmeal food supplemented with 50 mM metformin (Slack et al., 2012). Fifty larvae were transferred per vial (6 vials per genotype) and maintained at 25 +0.5^o^C.

**Adult lifespan analysis.** Lifespan was measured by maintaining newly eclosed adult *Drosophila* in cohorts of 25 flies per vial at 25 +0.5^o^C (Lin et al., 1998). Flies were transferred to fresh food vials every 2^nd^ day without anesthesia, and the number of dead individuals was recorded on each transfer. At least 100 flies per genotype were assayed across four replicate culture vials.

**SUPPLEMENTARY TABLES**

**Table S1: Reagents & Resources**

| ***Antibodies and stains*** | ***Source*** | ***Identifier*** |
| --- | --- | --- |
| Rabbit Anti-PH3 | Sigma-Aldrich | H0412 |
| Mouse Anti-Fas3 | DSHB | 7G10 |
| Rabbit Anti-p4E-BP | CST | 2855 |
| Rabbit Anti-ACC | CST | 3662S |
| Affinity-purified anti-Ilp2 (.55mg/ml) | Sangbin Park | (Park et al., 2014) |
| Alexa Fluor 488 Goat Anti-Rabbit | Invitrogen | A32731 |
| Alexa Fluor 633 Goat Anti-Rabbit | Invitrogen | A31210 |
| Alexa Fluor 488 Goat Anti-Mouse | Invitrogen | A32723 |
| Alexa Fluor 555 Goat Anti-Mouse | Invitrogen | A32727 |
| Alexa Fluor™ 488-Phalloidin | Invitrogen | A12379 |
| Alexa Fluor™ 555-Phalloidin | Invitrogen | A34055 |
| TO-PRO-3 iodide | Invitrogen | S33025 |
| Nile Red | Sigma-Aldrich | 19123 |
| DHE | Invitrogen | D11347 |
| PAS (Periodic acid & Schiff’s reagent) | S Ganesh, IITK | (Onkar et al., 2023) |
| F3 Carmoisine Red | Idacol-ROHA | CML63444 |
| ***Reagents and kits*** | ***Source*** | ***Identifier*** |
| Trehalase enzyme (Porcine Kidney) | Sigma-Aldrich | T8778-1UN |
| Glucose (HK) assay kit | Sigma-Aldrich | GAHK20-1KT |
| Free Glycerol Reagent | Sigma-Aldrich | F6428-40ML |
| Triglyceride Reagent | Sigma-Aldrich | T2449-10ML |
| Metformin Hydrochloride | Sigma-Aldrich | PHR1084-500MG |
| Schneider′s Insect Medium | Sigma-Aldrich | S9895 |
| p-Phenylenediamine hydrochloride | Sigma-Aldrich | P-6001-100G |
| TRI Reagent® | Sigma-Aldrich | T9424 |
| High-Capacity cDNA Reverse Transcription Kit | Applied Biosystems™ | 4368814 |
| PowerUp SYBR Green Master Mix | Applied Biosystems™ |  |
| Senescence cells histochemical staining kit | Sigma-Aldrich | CS0030 |
| CellEventTM Senescence Green Flow Cytometry Assay kit | Invitrogen | C10840 |
| Vectashield | Vector Labs | H-1200 |

**Table S2: *Drosophila* lines used in the study**

| ***Fly lines*** | ***Source*** | ***References*** |
| --- | --- | --- |
| *w1118* | BDSC (#5905) |  |
| *vg-Gal4* | Sean Carroll | (Williams et al., 1991) |
| *GMR-Gal4* | S. C. Lakhotia | (Li et al., 2012) |
| *Ov-Gal4 (Acp26A-Gal4)* | Ravi Ram | (Heifetz et al., 2000) |
| *UAS-GFP-nls* | BDSC (#4776) |  |
| *UAS-mRFP* | BDSC (#27392) |  |
| *UAS-Ras^G12V^* | Bruce Edgar | (Karim and Rubin, 1998) |
| *UAS-Yki3SA* | BDSC (#28817) |  |
| *UAS-Notch^intra^* | Marco Milan | (Go et al., 1998) |
| *UAS-scrib-IR* | BDSC (#29552) |  |
| *UAS-eiger-IR* | BDSC (#58993) |  |
| *UAS-impL2-IR* | BDSC (#64936) |  |
| *UAS-upd1-IR* | BDSC (#33680) |  |
| *UAS-upd2-IR* | BDSC (#33988) |  |
| *UAS-upd3-IR* | BDSC (#32859) |  |
| *UAS-hop-IR* | BDSC (#32966) |  |
| *UAS-impL2* | David Bilder | (Figueroa-Clarevega and Bilder, 2015) |
| *UAS-upd1* | Bruce Edgar | (Ren et al., 2010) |
| *UAS-upd2* | David Bilder | (Kim et al., 2021) |
| *UAS-upd3* | David Bilder | (Kim et al., 2021) |
| *UAS-eiger* | Jagat K Roy | (Igaki et al., 2002) |
| tGPH | BDSC (#8163) |  |
| STAT-10XGFP | BDSC (#26197) |  |
| ImpL2-EGFP-FLAG | BDSC (#59778) |  |
| Dilp2-mCherry | BDSC (#8163) |  |
| TRE-dsRed | BDSC (#59011) |  |

**Table S3: List of primers used in RT-PCR (qPCR)**

| **Genes** | **Forward primer (5'→3')** | **Reverse primer (5'→3')** |
| --- | --- | --- |
| *dILP2* | GGCCAGCTCCACAGTGAAGT | TCGCTGTCGGCACCGGGCAT |
| *dILP3* | CCAGGCCACCATGAAGTTGT | TTGAAGTTCACGGGGTCCAA |
| *dILP5* | TCCGCCCAGGCCGCAAACTC | TAATCGAATAGGCCCAAGGT |
| *upd1* | GTACCGCAGCCTAAACAGTAG | GCTTGGGCAGGAACTTGTA |
| *upd2* | CGGAACATCACGATGAGCGAAT | TCGGCAGGAACTTGTACTCG |
| *upd3* | TGGGAGAACACCTGCAATC | GCCCGTTTGGTTCTGTAGAT |
| *impL2* | CTCATGGGCTCCAACATTCA | CTTGGACGATCTCCTTGTTCTC |
| *eiger* | GTATGGGCTACCATGGAGATATG | CTGTATTGGTCACCGTCAAGA |
| *NFP* | TGCTCAGTCCAACTCCAACT | CACACGAGCAATAGCGATTC |
| *hugin* | TCTGCGTCTCTCTTTGTCGG | AAGGGAGAGGACACAGAATGA |
| *rpL23* | TGCCGGTACAAGAGGAGAAG | CGTTGTGGTTGTTGGTGATG |

**Table S4: Imaging and Software**

| ***Image acquisition*** | | | |
| --- | --- | --- | --- |
| Leica M205 FA stereomicroscope | | Leica microsystem | |
| Leica TCS SP5 confocal | | Leica microsystem | |
| Leica Stellaris confocal | | Leica microsystem | |
| Zeiss LSM710 Confocal Microscope | | Zeiss microsystem | |
| AXR Nikon Confocal Microscope | | NIS elements | |
| ***Software*** | | | |
| Leica LAS AF software | Leica microsystem | | www.leicamicrosystems.com/productes/microscope-software/ |
| FIJI | ImageJ | | imagej.net/software/fiji/ |
| Prism | GraphPad | | www.graphpad.com/ |
| Photoshop | Adobe CS-6 | | www.adobe.com |
| Illustrator | Adobe CC | | www.adobe.com |
| BioRender | Biorender Inc. | | biorender.com/ |

**SUPPLEMENTARY FIGURE LEGENDS**

**Fig. S1: *Ras^G12V^* triggers a localized metabolic reprogramming in the wing disc.**

**A)** *vg-Gal4* expression (using *vg-Gal4>UAS-GFPnls*) in the wing imaginal disc, haltere imaginal disc, and salivary glands. Nuclei stained for DAPI (blue).

**B-C)** p4e-BP (B), and ACC (C), both in red expression in the wing imaginal disc. *vg* domain marked with GFP (green), and nuclei stained for DAPI (blue).

**D)** SA-β-Gal activity in the wing (left panel) and salivary glands (right panel).

**E)** qPCR analysis of different cytokines expression from the salivary glands. (N=3 biological replicates, *****p<0.0001* and ***p<0.01*, Data are presented as mean ± SEM).

Scale bars: 50µm (A, B, and C), and 200µm (D).

**Fig S2: *Ras^G12V^*-expressing adults display cachexia-like phenotype.**

**A)** Comparison of larval to adult fraction in wild type and *Ras^G12V^* expressing adults (N=100 larvae per genotype, *****p<0.0001*).

**B)** Comparison of adult lifespan of respective genotypes (N=100 adults observed per genotype, *****p<0.0001*).

**C)** Adults (first panel), abdominal fat body (Nile red, second panel), abdominal muscles (Actin, third panel), and adult ovary (Actin (green) and DAPI (blue)) of respective genotypes (N=10 for each genotype).

Scale bars: 1mm (C, first panel), 50µm (C, rest of the panels).

**Fig. S3: Senescence activation in fat body is independent of the JAK-STAT pathway, and Upd1 works as a senescence relay cytokine.
A)** STAT-GFP expression in the wing disc (upper panel) and fat body (lower panel) in wild-type compared to *Ras^G12V^*-expressing larvae.

**B)** SA-β-Gal activity (enzymatic) in the male accessory glands (MAG) upon ectopic expression of different cytokines.

**C)** SA-β-Gal activity in the adult fat body upon ectopic expression of different cytokines from MAG.

Scale bars: 50µm (A), 200µm (B), and 25µm (C).

**Fig. S4: Senescence induction and its relay, a cell-type-oncogene-code dependent mechanism.
A)** SA-β-Gal activity (green) in the wing disc (upper panel) and the fat body (bottom panel) upon expression of *N^intra^*, *Yki^3SA^*, and *scrib*-*IR*, *Ras^G12V^* in the wing disc. RFP marks the *vg* domain (*vg-Gal4>UAS-RFP*).

**B)** SA-β-Gal activity (green) in the eye disc (upper panel) and the fat body (bottom panel) upon expression of *Ras^G12V^*, *Yki^3SA^*, *N^intra^*, and *scrib*-*IR*, *Ras^G12V^* in the eye disc. RFP marks the *vg* domain (*vg-Gal4>UAS-RFP*), and nuclei are marked with DAPI.

**C)** SA-β-Gal activity (green) in the MAG (upper panel) and the fat body (bottom panel) upon expression of *N^intra^*, *Yki^3SA^*, and *scrib*-*IR*, *Ras^G12V^* in the wing disc.

**D)** SA-β-Gal activity (green) in the larval muscles upon *Ras^G12V^* and *scrib*-*IR*, *Ras^G12V^* expression in the wing disc.

Scale bar: 50µm.

**Fig. S5: Epithelial *Ras^G12V^* elevates JAK-STAT pathway activity in the larval brain.**

**A)** Larval brains expressing dILP2-mCherry and STAT-10XGFP in wild type and upon *Ras^G12V^* expression.

Scale bar: 25µm.

**Figure S6: Metformin suppresses tumor growth.**

**A)** Adult eclosion fraction upon genetic knockdown of *upd1* in the *Ras^G12V^*-expressing larvae as compared to *Ras^G12V^*-expressing larvae alone and wild type (N=10 larval to adult transition analyzed, *****p<0.0001,* Data are presented as mean ± SEM).

**B-C)** SA-β-Gal activity (green, B) in the neoplastic *scrib-IR*; *R­as^G12V^* tumor and their size quantification (C) upon metformin feeding.

Scale bar: 50µm.
