## Supplementary figures and images for "*Ras^G12V^* Oncogene-Induced Epithelial Senescence and Its Relay Promotes Host Metabolic Syndrome in *Drosophila*"

### Fig. S1

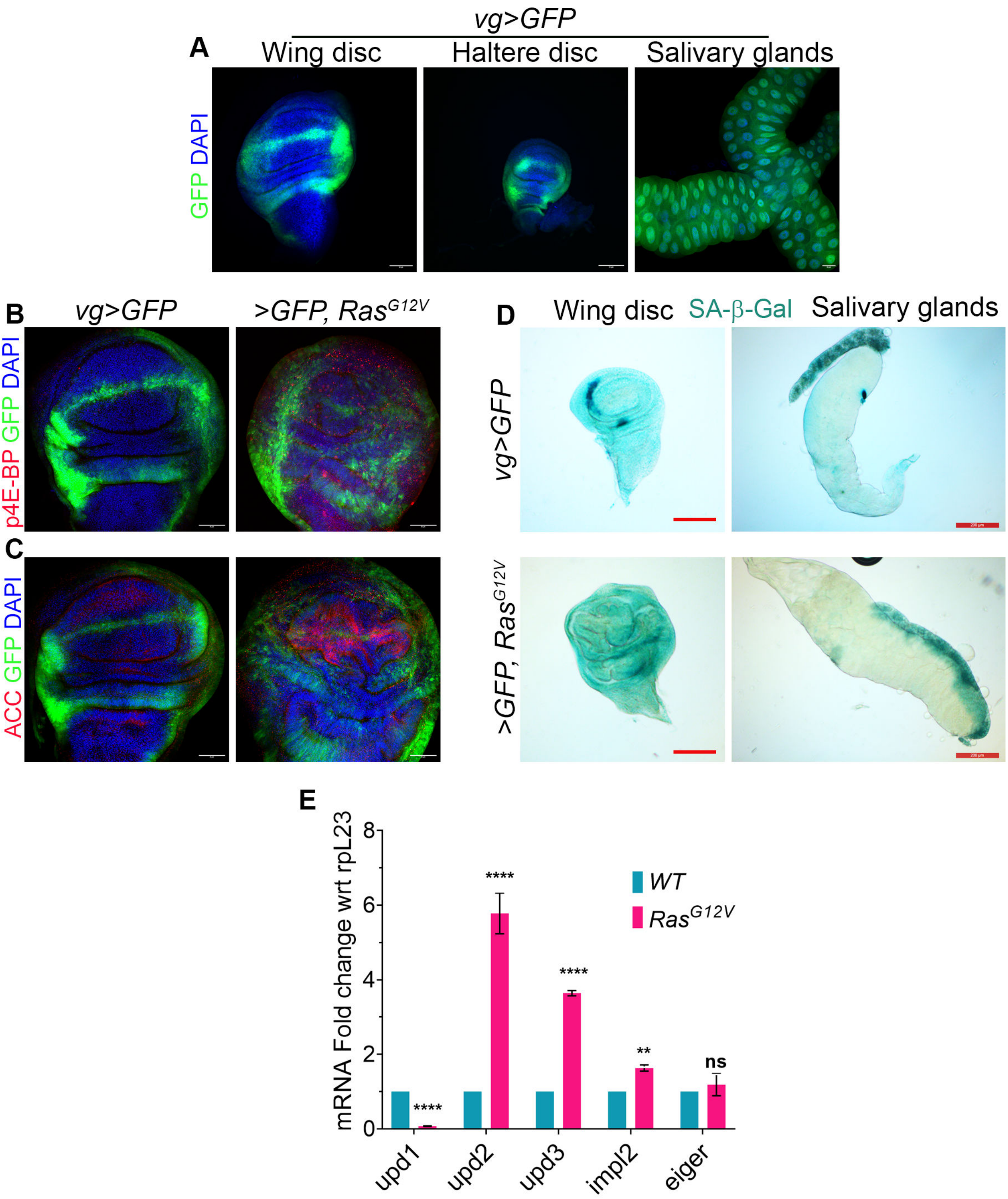

### Fig. S2

**Fig. S2**

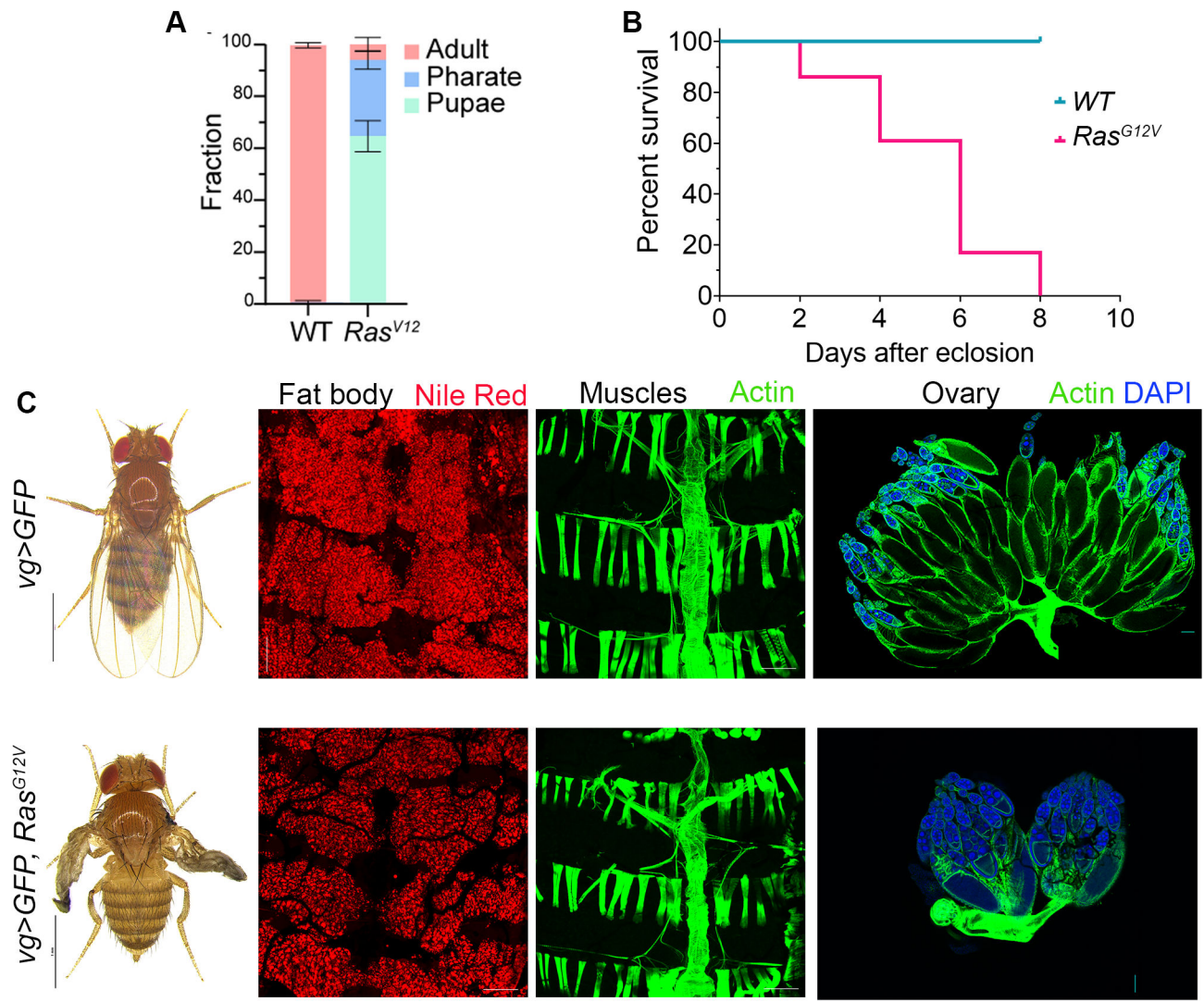

### Fig. S3

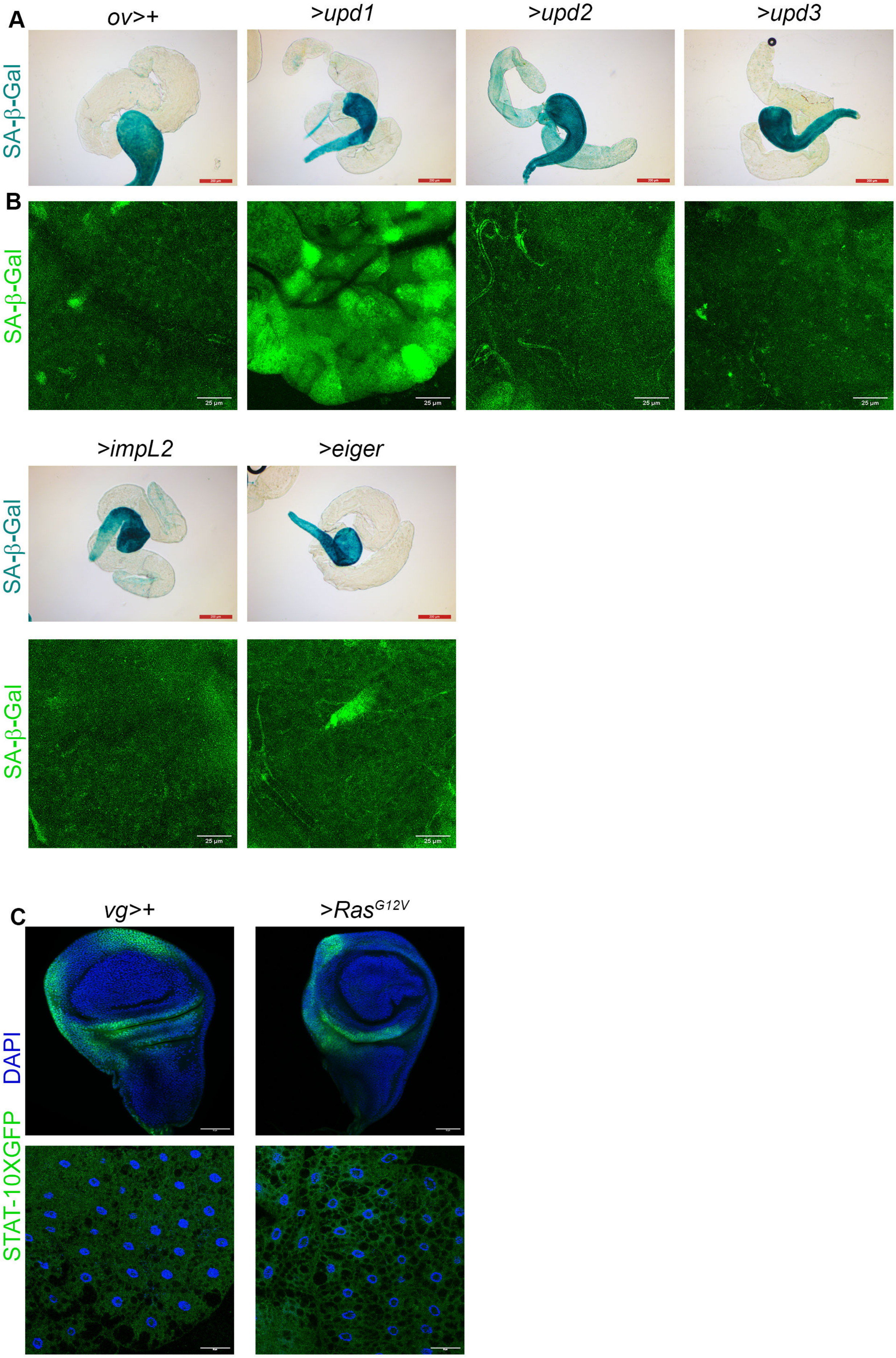

### Fig. S4

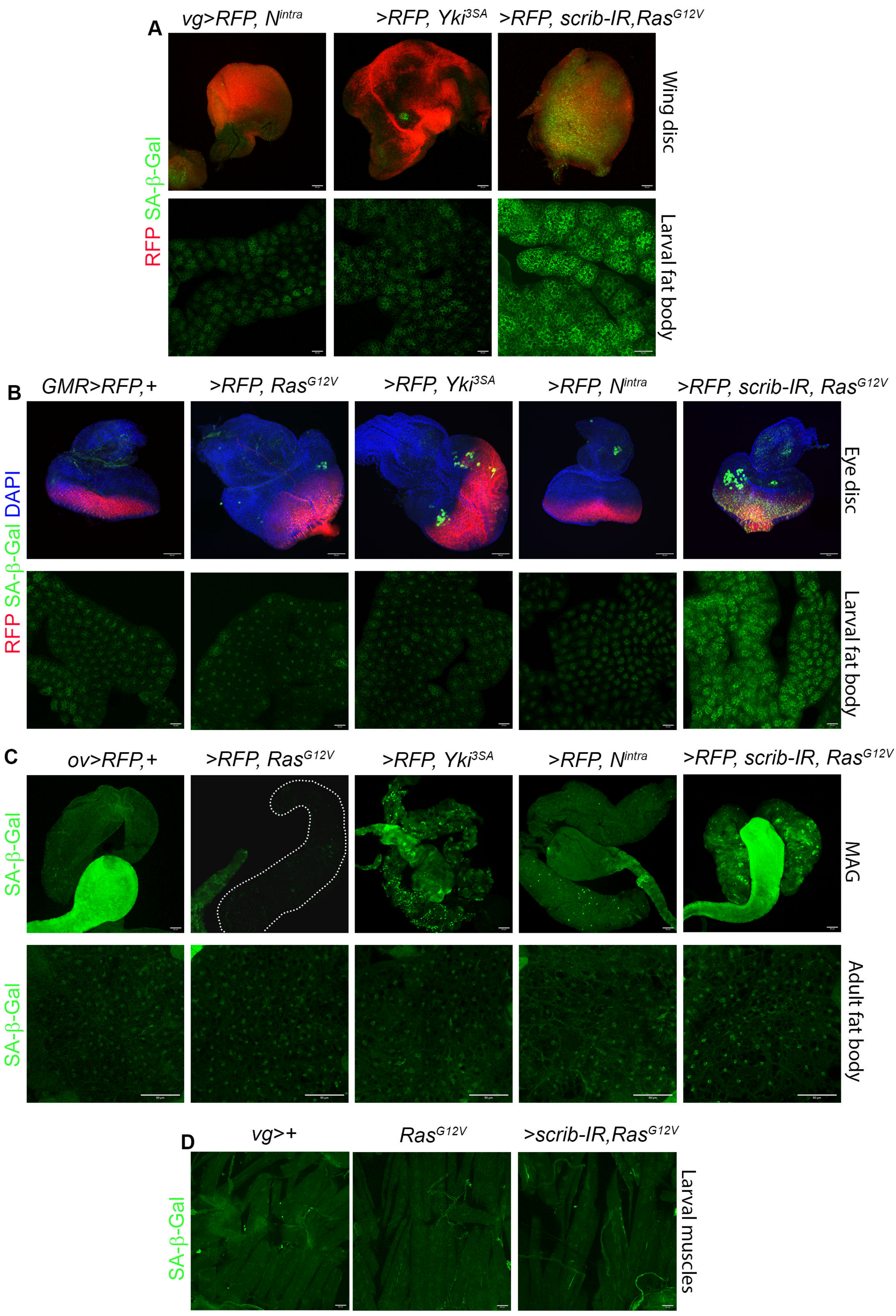

### Fig. S5

#### Fig. S5

**A**

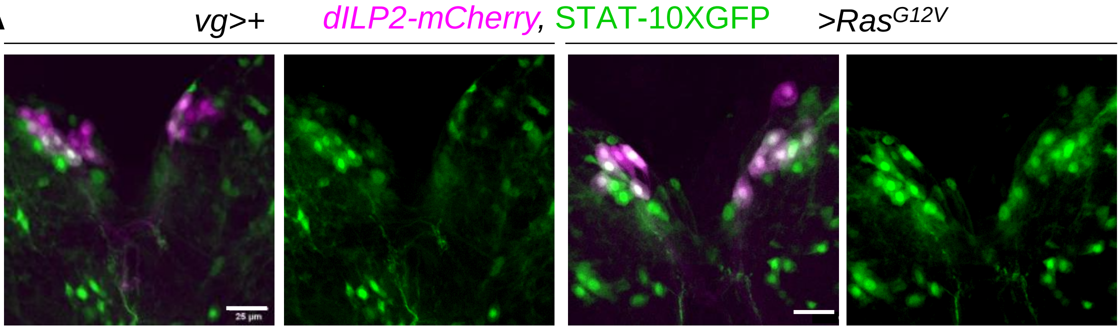

### Fig. S6

#### Supplementary Figure 6

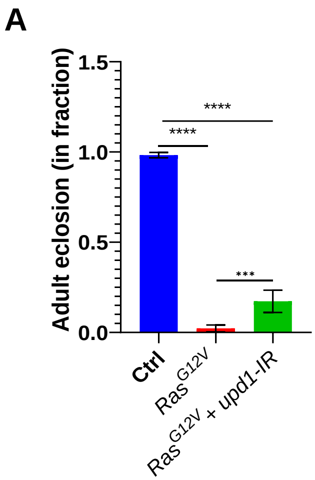


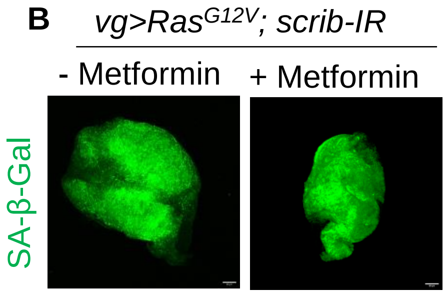


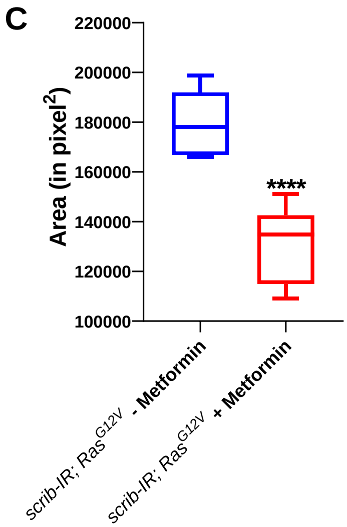
